# Addiction of mesenchymal phenotypes on the FGF/FGFR axis in oral squamous cell carcinoma cells

**DOI:** 10.1101/638387

**Authors:** Asami Hotta Osada, Kaori Endo, Yujiro Kimura, Kei Sakamoto, Ryosuke Nakamura, Kaname Sakamoto, Koichiro Ueki, Kunio Yoshizawa, Keiji Miyazawa, Masao Saitoh

**Affiliations:** Department of Biochemistry, Interdisciplinary Graduate School of Medicine, University of Yamanashi, Chuo, Yamanashi; Department of Oral and Maxillofacial Surgery, Interdisciplinary Graduate School of Medicine, University of Yamanashi, Chuo, Yamanashi; Department of Center for Medical Education and Sciences, Interdisciplinary Graduate School of Medicine, University of Yamanashi, Chuo, Yamanashi; Department of Oral Pathology, Graduate School of Medical and Dental Sciences, Tokyo Medical and Dental University, Yushima, Bunkyo-ku, Tokyo; Department of Oral Surgery, Kofu Municipal Hospital, Kofu, Yamanashi, Japan

**Author notes:** Corresponding author: Masao Saitoh, Department of Biological Chemistry, Center for Medical Education and Sciences, Interdisciplinary Graduate School of Medicine, University of Yamanashi, 1110 Shimokato, Chuo, Yamanashi 409-3898, Japan.

## Abstract

The epithelial–mesenchymal transition (EMT) is a crucial morphological event that occurs during epithelial tumor progression. ZEB1/2 are EMT transcription factors that are positively correlated with EMT phenotypes and breast cancer aggressiveness. ZEB1/2 regulate the alternative splicing and hence isoform switching of fibroblast growth factor receptors (FGFRs) by repressing the epithelial splicing regulatory proteins, ESRP1 and ESRP2. Here, we show that the mesenchymal-like phenotypes of oral squamous cell carcinoma (OSCC) cells are dependent on autocrine FGF–FGFR signaling. Mesenchymal-like OSCC cells express low levels of ESRP1/2 and high levels of ZEB1/2, resulting in constitutive expression of the IIIc-isoform of FGFR, FGFR(IIIc). By contrast, epithelial-like OSCC cells showed opposite expression profiles for these proteins and constitutive expression of the IIIb-isoform of FGFR2, FGFR2(IIIb). Importantly, ERK was constitutively phosphorylated through FGFR1(IIIc), which was activated by factors secreted autonomously by mesenchymal-like OSCC cells and involved in sustained high-level expression of ZEB1. Antagonizing FGFR1 with either an inhibitor or siRNAs considerably repressed ZEB1 expression and restored epithelial-like traits. Therefore, autocrine FGF–FGFR(IIIc) signaling appears to be responsible for sustaining ZEB1/2 at high levels and the EMT phenotype in OSCC cells.

## Introduction

Oral tongue squamous cell carcinoma (OSCC) is one of the most common malignancies in head and neck cancers[1]. Most patients on their first visit to hospitals are diagnosed with locoregional initial symptoms of the disease. After various treatments, including surgical operation, chemotherapy, radiotherapy, or combination, the five-year survival rate remains less than 50% due to its aggressive invasiveness and resistance to treatments[2]. Thus, the development of diagnostic and therapeutic strategy would be of significant benefit to the development of successful therapies.

The process of cancer cell invasion involves the loss of cell–cell interactions along with the acquisition of motility, and is partly associated with the epithelial–mesenchymal transition (EMT)[3]. EMT involves dramatic cellular changes in which epithelial cells loosen their attachments to neighboring cells, lose their apico-basal polarity, become elongated, and display increased motility. EMT therefore forms the first step of the invasion–metastasis cascade. After invading through basement membranes and blood/lymphatic vessel walls, cells undergoing EMT survive in the bloodstream as circulating tumor cells, and, lastly, extravasate into distant organs. Upon arriving at distant metastasized tissues, the cancer cells undergo a reversion process, the mesenchymal–epithelial transition (MET)[4, 5]. The EMT process is regulated by several transcription factors known as EMT transcription factors (EMT-TFs), including the δEF1 family of two-handed zinc-finger factors (ZEB1 [Zinc-finger E-box binding homeobox 1]/δEF1 [δ-crystallin/E2-box factor 1] and ZEB2/SIP1 [Smad-interacting protein1]), the Snai1 family (Snail, Slug, and Smuc), and basic helix-loop-helix factors (Twist and E12/E47). Among these, the levels of ZEB1/2 in particular correlate positively with EMT phenotypes and the aggressiveness of breast cancer cells[6, 7]. It remains unclear, however, why expression of ZEB1/2 is sustained at high levels in aggressive cancer cells.

The alternative splicing machinery is also involved in regulating EMT[4]. Epithelial splicing regulatory proteins (ESRPs) 1 and 2, also known as RNA-recognition motif–containing proteins Rbm35a and Rbm35b, respectively, induce the switching of alternative splicing of transcripts, such as FGF receptors (FGFRs), CD44, Rac1, p120 catenin, and Mena. ZEB1/2 are preferentially recruited to the promoter region of *ESRP1*, and suppress the transcription of *ESRP1* during EMT[8, 9]. Despite the similar primary structures of the ESRP1 and ESRP2 proteins, the functions of the two proteins differ slightly in OSCC cells[10].

The *FGFR* genes encode four functional receptors (FGFR1–4) with three extracellular immunoglobulin-like domains, namely, Ig-I, Ig-II, and Ig-III. The Ig-III domain is regulated by alternative splicing, which produces either the IIIb isoforms, FGFR1(IIIb)–FGFR3(IIIb), or the IIIc isoforms, FGFR1(IIIc)–FGFR3(IIIc), which have distinct FGF binding specificities[11]. Mesenchymal cells expressing the IIIc-isoform respond to FGF2, also known as basic FGF, and FGF4. By contrast, epithelial cells generally expressing the IIIb isoform consequently respond to FGF7, also known as keratinocyte growth factor (KGF), and FGF10. In fact, cancer cells with low expression of ESRP1/2 and high expression of ZEB1/2, are associated with aggressive behavior and poor prognosis, and express only the IIIc isoforms. Conversely, cells that express low levels of ZEB1/2 and high levels of ESRP1/2 are associated with favorable prognoses, and exhibit constitutive expression of the IIIb isoforms[6].

In this study, we determined the EMT phenotypes of OSCC cells and found that FGFR2-IIIb was ubiquitously expressed in epithelial-like OSCC cells. Among various OSCC cells, we determined that TSU and HOC313 cells exhibited mesenchymal-like phenotypes with high motility. In addition, we found that TSU and HOC313 cells exhibited high levels of phosphorylated extracellular signal-regulated kinase 1/2 (ERK1/2), and expressed low levels of ESRP1/2 along with high levels of ZEB1/2 levels, resulting in constitutive expression of only FGFR1(IIIc). The FGFR1(IIIc) isoform is apparently activated by soluble factors secreted autonomously by these cells and is needed to sustain high-level expression of ZEB1/2. When we antagonized FGFR1 by either using an inhibitor or specific siRNAs, resulting in the inactivation of ERK1/2 and repression of ZEB1/ZEB2, we observed partial phenotypic changes to epithelial traits. Therefore, sustained high-level expression of ZEB1/2 mediated by the FGFR1c-ERK pathway may maintain the mesenchymal-like phenotypes of OSCC cells.

## Materials and methods

### Cell Culture

Human OSCC, TSU, HOC313, OBC-01, OSC-19, OSC-20, and OTC-04 cells were gifts from Dr. E. Yamamoto and Dr. S. Kawashiri[12]. HSC-2, HSC-3, and HSC-4 were gifts from Dr. F. Momose and Dr. H. Ichijo[13, 14]. SAS and Ca9-22 cells were described previously[15]. All cells were cultured in DMEM (Nacalai Tesque, Kyoto, Japan) supplemented with 4.5 g/L glucose, 10% FBS, 50 U/mL penicillin, and 50 μg/mL streptomycin at 37°C under a 5% CO_2_ atmosphere.

### Reagents and Antibodies

Recombinant human TGF-β, FGF basic (FGF2), and FGF7 were obtained from R&D Systems (Minneapolis, MN). Rabbit monoclonal anti–phospho-ERK1/2 antibody was from Cell Signaling (Danvers, MA). Rabbit polyclonal anti-ZEB1 and anti-ZEB2 antibodies were obtained from Novus Biologicals (Littleton, CO). Mouse monoclonal anti–E-cadherin and anti–α-tubulin antibodies were from BD Biosciences (Lexington, KY) and Sigma-Aldrich (St. Louis, MO), respectively. SU5402 and U0126 were purchased from Calbiochem (Darmstadt, Germany) and Promega (Madison, WI), respectively. AP24534 was from Selleck Chemicals (Houston, TX).

### Immunoblotting and immunofluorescence

The procedures used for immunoblotting and immunofluorescence assays were previously described[16]. Briefly, cells were lysed in lysis buffer (20 mM Tris-HCl [pH7.5], 150 mM NaCl, 1% Nonidet P-40, 5 mM EDTA, 1 mM EGTA, and protease and phosphatase inhibitors). Protein concentration was measured using BCA protein assay reagent (Thermo Fisher Scientific, Waltham, MA). Harvested proteins separated by SDS-PAGE were transferred on to polyvinylidene difluoride membranes, followed by immunodetection with the ECL western blotting system (GE Healthcare, Piscataway, NJ) on a Luminescent Image Analyzer (LAS400, Fujifilm, Tokyo, Japan).

### Quantitative Real-time PCR (qRT-PCR)

Total RNA was extracted using the RNeasy mini kit (Qiagen, Venlo, Netherlands) and cDNAs were synthesized using the PrimeScript First Strand cDNA synthesis kit (Takara-Bio, Kusatsu, Japan). qRT-PCR analyses were performed using Power SYBR Green PCR Master Mix (Applied Biosystems, Foster City, CA). The relative expression level of each mRNA was normalized against level of *GAPDH* mRNA. The primers used were described previously[6], except the primers specific for human *FGFR1*, *FGFR2*, and *FGF2*.

human *FGFR1*: forward, 5’-TGAGTACGGCAGCATCAACCAC-3’; reverse, 5’-ACTGTTTTGTTGGCGGGCAAC-3’

human *FGFR2*: forward, 5’-TGTGCACAAGCTGACCAAACG-3’; reverse, 5’-AGGCGTGTTGTTATCCTCACCAG-3’

human *FGF2*: forward, 5’-AACCTGCAGACTGCTTTTTGCC; reverse, 5’-ACGTGAGAGCAGAGCATGTGAG

### RNA interference

Transfection of siRNAs was performed in six-well tissue culture plates using Lipofectamine RNAiMAX transfection reagent (Invitrogen). The final concentration of siRNA was 10 nM. The stealth RNAi siRNA against either human *FGFR1* or mouse *Fgfr1* were purchased from Thermo Fisher Scientific.

### Cell proliferation assay

Cells were seeded on six-well plates, then trypsinized and counted by hemocytometer. Twenty-four hours after transfection with the siRNAs, cells were seeded in triplicate in 24-well tissue culture plates. Twenty-four hours after seeding, cell count assays were carried out using Cell Count Reagent SF (Nacalai Tesque).

### Invasion assay

Boyden chamber migration assays were conducted using transparent PET membrane 24-well 8.0 μm pore size cell culture inserts (BD Falcon, Franklin Lakes, NJ) coated with collagen type I-C (Nitta Gelatin, Osaka, Japan). After cells were seeded in triplicate on the inserts, cells that had not invaded the lower surfaces of the filters were removed from the upper faces of the filters using cotton swabs. Cells that invaded into the lower surfaces of the filters were fixed in acetone:methanol (1:1) and stained with Trypan Blue. Invasion was quantitated by visually counting photographed cells. Cell numbers were evaluated by statistical analysis.

### Statistical analyses

Data are presented as means ± SD. Statistical analyses were performed using Student’s *t*-test between any two groups.

## Results

### Evaluation of EMT phenotypes in human OSCC cell lines

To determine the EMT phenotypes of human OSCC cells, we investigated OSCC cell lines by immunoblot and qRT-PCR analyses. Among various OSCC cells, TSU and HOC313 cells expressed high levels of vimentin and low levels of E-cadherin, while other OSCC cells expressed high levels of E-cadherin and low levels of vimentin (Figs. 1A, 1B, and 1C). OTC-04 cells exhibited a hybrid phenotype, expressing both E-cadherin and vimentin. (Fig. 1A). HSC-4 cells exhibited a cobblestone-like shape, whereas TSU and HOC313 cells showed a spindle-like shape with greater motility compared to other epithelial-like OSCC cells (Figs. 1D and 1E), suggesting that both TSU and HOC313 cells show mesenchymal-like trails, whereas other OSCC cells exhibit epithelial-like characteristics.

**Figure 1.**
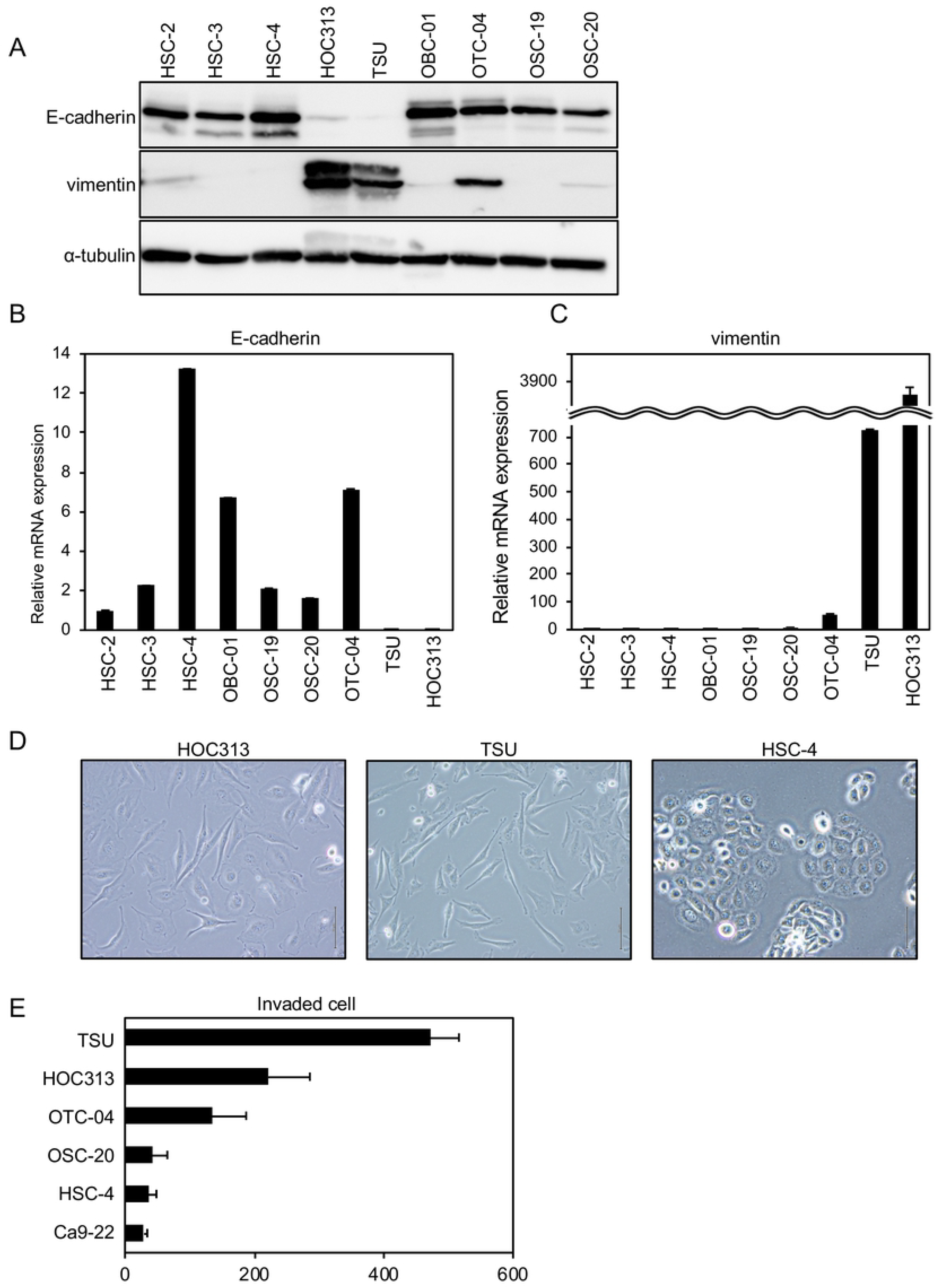
E-cadherin and vimentin expression profiles, and motility of various OSCC cells. **(A, B, C)** Expression levels of E-cadherin and vimentin were determined by immunoblotting (A) and qRT-PCR (B and C). For immunoblotting, α-tubulin levels were monitored as a loading control (A). **(D)** Representative images of showing the morphology of TSU, HOC313, and HSC-4 cells. **(E)** Motility of various OSCC cell lines was determined by transwell assays. Each value represents the mean ± SD of triplicate determinations from a representative experiment. Similar results were obtained from at least three independent experiments (B, C, E)

### Determination of ZEB1/2 expression in OSCC cells

We previously reported that the expression of ZEB1/2 is positively correlated with EMT phenotypes of breast cancer cell lines[6, 7]. In breast cancer, cells with high levels of ZEB1/2 and low levels of ESRP1/2 and E-cadherin are categorized into the “basal-like” subtype of breast cancer with aggressive behavior and poor prognosis[6, 17]. By contrast, cells that express low levels of ZEB1/2 along with high levels of ESRP1/2 and E-cadherin were categorized into the “luminal” subtype of breast cancer with relatively good prognosis[6, 17]. Similar to the basal-like subtype, mesenchymal-like OSCC, TSU and HOC313, cells exhibited high levels of *ZEB1* and *ZEB2* mRNA and low levels of *ESRP1* and *ESRP2* mRNA (Figs. 2A and S1A). On the other hand, the other epithelial-like OSCC cells showed the opposite expression profiles for these mRNAs (Figs. 2A and S1A). OSCC tissues expressing high levels of *ZEB1* also showed high levels of *ZEB2* (Fig. S1B). However, the expression levels of neither Snail nor Slug were faithfully correlated with the epithelial-like phenotypes of OSCC cells used in this study (Fig. S1C).

**Figure 2.**
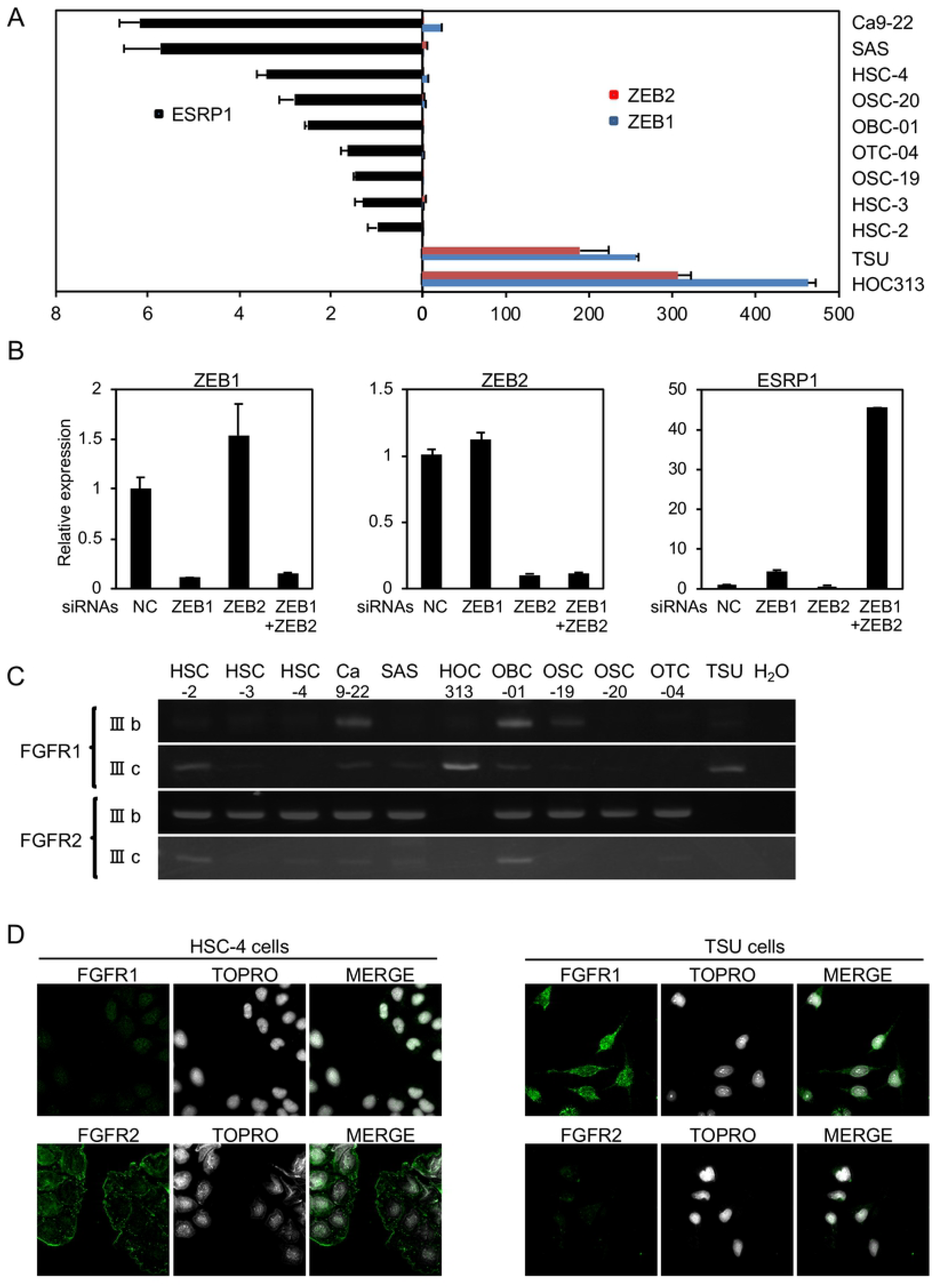
*ESRP1*, *ZEB1*, and *ZEB2* expression profiles in OSCC cells. **(A)** mRNA levels of the expression of *ESRP1*, *ZEB1*, and *ZEB2* were determined by qRT-PCR. Each value was normalized to the level of *GAPDH* in the same sample. **(B)** TSU cells were transfected with siRNAs against *ZEB1*, *ZEB2*, or both, and *ESRP1* mRNA level was determined by qRT-PCR. NC, non-specific control siRNA. Each value represents the mean ± SD of triplicate determinations from a representative experiment. Similar results were obtained in at least three independent experiments. **(C)** The expression of FGFR isoforms in OSCC cells was determined by conventional RT-PCR. **(D)** Subcellular localization of endogenous FGFR1 and FGFR2 proteins was determined by anti–FGFR1 and –FGFR2 antibodies, respectively, in HSC-4 and TSU cells.

We also previously reported that ZEB1/2 are preferentially recruited to the promoter region of *ESRP1* where they suppress the transcription of *ESRP1* in breast cancer cells[6]. When both *ZEB1* and *ZEB2* are simultaneously knocked down with their specific siRNAs, *ESRP1* expression was dramatically upregulated in HOC313 cells, whereas *ESRP2* was not (Fig. 2B and data not shown); this phenotype did not occur when either ZEB1 or ZEB2 was knocked down alone. Since ESRP1, rather than ESRP2, causes the alternative splicing-mediated isoform switching between *RAC1b* transcripts in OSCC cells[10], we determined the relative abundance of alternative splicing variants of *FGFR1* and *FGFR2*. Based on qRT-PCR, we were unable to reliably detect *FGFR3* and *FGFR4* expression in the cells, as indicated by qRT-PCR (data not shown). Hence, we were only able to investigate the alternative splicing of *FGFR1*/*2*.

Interestingly, all epithelial-like OSCC cells expressed FGFR2(IIIb), which was not detected in either TSU or HOC313 cells. By contrast, TSU and HOC313 cells expressed only FGFR1(IIIc) (Fig. 2C). Indeed, analyses using the TCGA dataset indicated a positive correlation between ZEB1/2 and FGFR1, and negative correlation between ZEB1/2 and FGFR2 (Figs. S2A and S2B). HSC-2 cells expressed both the IIIb and IIIc isoforms of FGFRs (Fig. 2C), probably due to low expression of both *ESRP1* and *ESRP2*, whereas OBC-01 cells also expressed both isoforms with moderate expression of *ESRP1*/*2* (Figs. 2A, and S1A). Although OTC-04 cells exhibited a hybrid phenotype, only FGFR2(IIIb) was observed (Fig. 2C). In HSC-4 cells, FGFR2 was localized to the plasma membrane while only negligible amounts of FGFR1 were detected. TSU cells exhibited almost no detectable FGFR2 while the intracellular localization of FGFR1 was diffused (Fig. 2D).

### Constitutive activation of ERK1/2 in mesenchymal-like OSCC cells

FGF2 and FGF4 bind preferentially to the IIIc-isoform, whereas FGF7 and FGF10 bind exclusively to the IIIb-isoform[11]. In HSC-4 and OTC-04 cells, FGF7 induced the phosphorylation of ERK1/2, but FGF2 did not due to the lack of IIIc-isoform expression in the cells (Fig. 3A). Interestingly, ERK1/2 phosphorylation in mesenchymal-like TSU and HOC313 cells was detected even under serum-free culture conditions (Fig. 3B). Although FGF2 induced only slight phosphorylation of ERK1/2 in cells that express FGFR1(IIIc) (Fig. 3B), treatment with the FGFR1 inhibitor, SU5402, almost completely inhibited phosphorylation of ERK1/2, suggesting the involvement of autocrine factors that activate FGFR1(IIIc). To test this possibility, conditioned medium from TSU cells was added to mouse mammary epithelial NMuMG cells pretreated with TGF-β to express Fgfr1(IIIc)[16]. The conditioned medium from TSU cells, but not HSC-4 cells, caused a slight increase in ERK1/2 phosphorylation, while either pretreatment with SU5402 or transfection with mouse *Fgrf1* siRNA into TGF-β-treated NMuMG cells repressed it (Fig. 3C, and data not shown).

**Figure 3.**
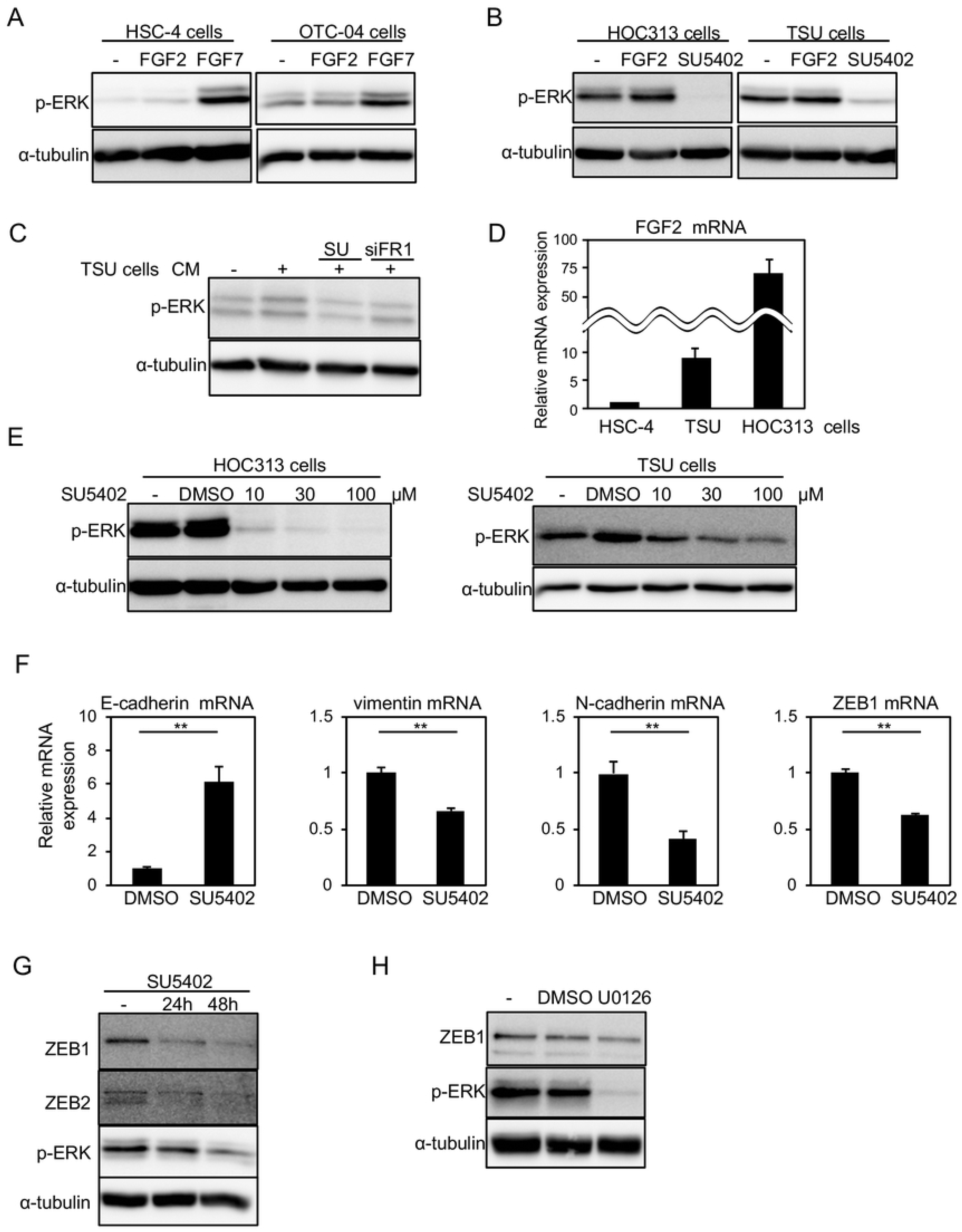
SU5402, FGFR1 inhibitor, affects the EMT transcription factors. **(A, B)** ERK1/2 phosphorylation (p-ERK) was determined by immunoblotting in HSC-4 and OTC-04 treated for 30 min with 30 ng/ml FGF2 or 30 ng/ml FGF7 in the presence of 10% FBS (A) and in TSU and HOC313 cells treated for 1 h with 30 ng/ml FGF2 or 30 μM SU5402 in the absence of FBS. F2, FGF2; F7, FGF7; SU, SU5402. **(C)** We have previously reported that, after treatment with TGF-β, NMuMG cells underwent EMT with the IIIc-isoform of FGFR1[16]. After NMuMG cells pretreated with TGF-β were transfected with mouse *Fgfr1* siRNA or treated with SU5402, the cells were further incubated in culture medium (CM) from TSU cells. SU, SU5402; siFR1, siRNA against mouse *Fgfr1*. **(D)** *FGF2* mRNA levels were determined by qRT-PCR analyses. *FGF2* mRNA levels in HSC-4 cells were indicated as 1. Each value represents the mean ± SD of triplicate determinations from a representative experiment. Similar results were obtained in at least three independent experiments. **(E)** ERK1/2 phosphorylation (p-ERK) in TSU and HOC313 cells were monitored in the presence of the indicated concentration of SU5402 for 1 h under serum-free culture conditions, followed by immunoblot analysis. **(F, G)** Expression of the indicated genes in TSU cells under serum-free culture conditions was determined by qRT-PCR (D) and immunoblot (F) analyses, following treatment with 10 μM SU5402. Each value represents the mean ± SD of triplicate determinations from a representative experiment. Similar results were obtained from at least three independent experiments. p values were determined by Student’s t-test. **p < 0.01. **(H)** TSU cells treated with 10 μM U0126 in the absence of FBS were subjected to immunoblotting with the indicated antibodies. α-tubulin was used as a loading control (A, B, C, E, G, H).

Indeed, *FGF2* mRNA was highly expressed in mesenchymal-like TSU and HOC313 cells (Fig. 3D). SU5402 inhibited phosphorylation of ERK1/2 in a dose-dependent manner in HOC313 and TSU cells (Fig. 3E). In addition, the representative epithelial marker, E-cadherin, was upregulated by SU5402, whereas the representative mesenchymal markers, N-cadherin and vimentin, as well as ZEB1/2 were suppressed at both the mRNA and protein levels (Figs. 3F and 3G, and data not shown). These findings suggest that mesenchymal-like OSCC cells autonomously secrete factors, including FGF2, to activate FGFR(IIIc), and that blocking FGFR1 signaling regulates the expression of EMT markers. Importantly, ZEB1 was retained at high levels in mesenchymal-like TSU and HOC313 cells, which was repressed by FGFR1 inhibitor. When ERK was also inactivated by the MEK inhibitor, U0126, ZEB1 was downregulated (Fig. 3H), strongly suggesting that high-level expression of ZEB1 is sustained by constitutive activation of ERK1/2 mediated by FGFR(IIIc), which is activated by soluble factors secreted autonomously by the cells.

### Roles of *FGFR1* siRNAs in mesenchymal-like TSU cells

In addition to SU5402, to elucidate the functions of FGFR1 in mesenchymal-like TSU cells, we used siRNAs against human *FGFR1*. Three kinds of *FGFR1*-targeted siRNAs effectively knocked down endogenous *FGFR1*, as determined by conventional RT-PCR analyses (Fig. 4A). ERK1/2 phosphorylation and ZEB1 expression in TSU cells under serum-free culture condition were reduced by *FGFR1* siRNAs (Figs. 4B and 4C). The repressive effects of *FGFR1* siRNAs on ZEB1 levels were also observed in the basal-like breast cancer, Hs-578T, and MDA-MB231 cells, which expressed FGFR1(IIIc) isoform (Fig. S3A)[6]. In addition, cell morphology was slightly altered from a spindle shape to a cobblestone-like shape (Fig. 4D). Immunofluorescence analyses indicated that E-cadherin was upregulated and localized to the plasma membrane in some TSU cells (Fig. 4E).

**Figure 4.**
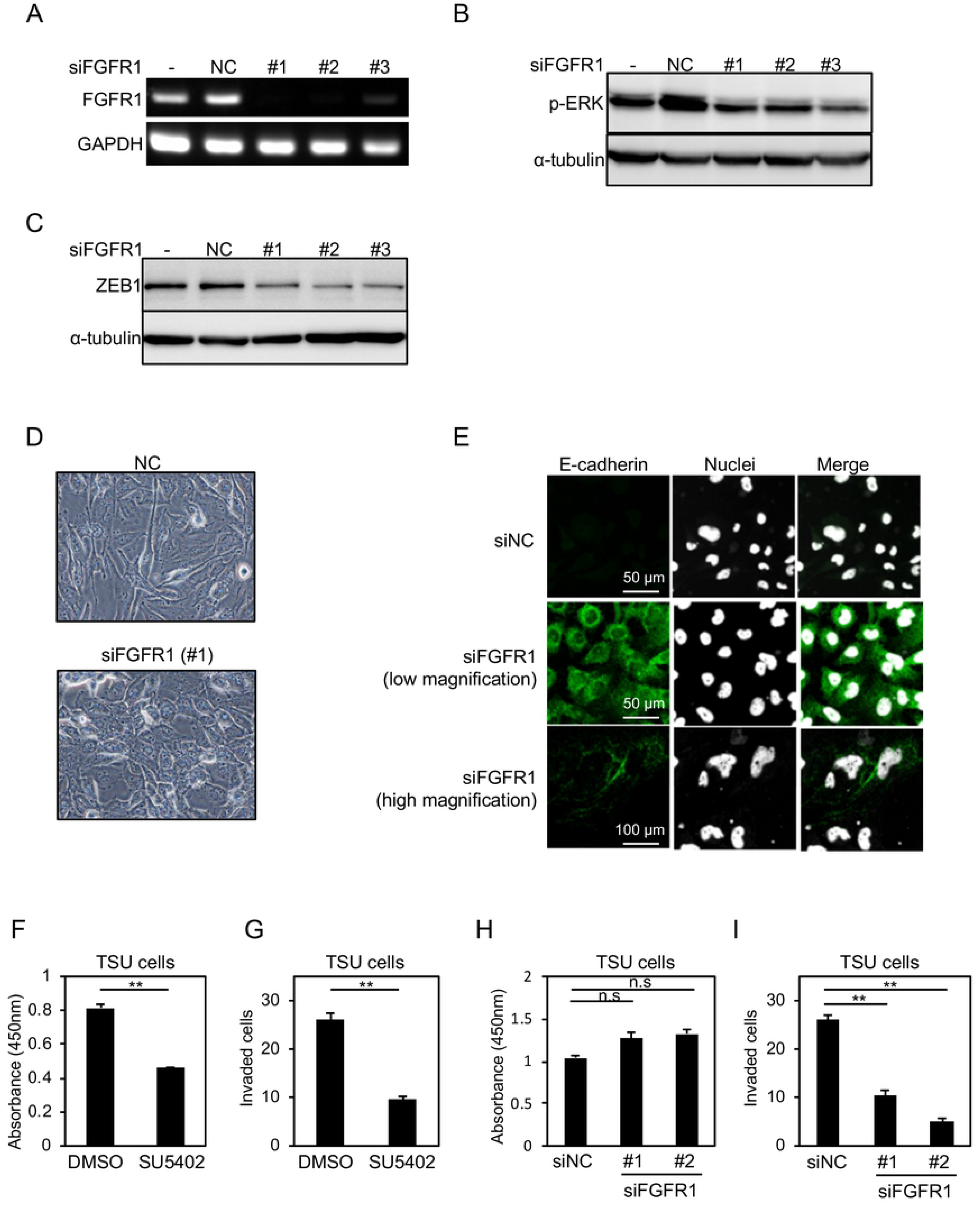
*FGFR1* siRNAs attenuate the malignant phenotypes of cancer cells. **(A)** mRNA from TSU cells transfected with siRNAs against *FGFR1* (si*FGFR1*) were subjected to conventional RT-PCR to determine the levels of endogenous *FGFR1*. **(B, C, D)** After transfection with si*FGFR1* in TSU cells, phosphorylation of ERK1/2 (p-ERK) (B), ZEB1 levels (C), and cell morphology (D) were determined under serum-free culture conditions. **(E)** TSU cells transfected with si*FGFR1* were subjected to immunofluorescence analyses. Low magnification, 40×; high magnification, 100×. **(F, G, H, I)** After either treatment with SU5402 or transfection with si*FGFR1* in TSU cells, the number of cells (F, H) and invasive properties (G, I) were determined under serum-free culture conditions. NC, negative control siRNA. Each value represents the mean ± SD of triplicate determinations from a representative experiment. Similar results were obtained from at least three independent experiments. p values were determined by Student’s t-test. *p < 0.01; n.s., not significant.

Invasion properties were inhibited by SU5402, which was accompanied with the reduced cell number (Figs. 4F and 4G). Another FGFR1 inhibitor, AP24534, also suppressed the motility of TSU cells (Fig. S3B). By contrast, siRNAs against *FGFR1* inhibited invasive properties without drastically affecting the number of cells (Figs. 4H and 4I). Taken together, FGFR(IIIc) isoforms, which are predominantly expressed in mesenchymal-like OSCC cells, would be constitutively activated by factors secreted autonomously by the cells and sustain ZEB1 expression at high levels through the activating ERK pathway, leading to maintaining EMT phenotypes.

## Discussion

In the study we found that, similar to the basal-like subtype of breast cancer, OSCC cells with high levels of ZEB1/2 and low levels of E-cadherin and ESRP1/2 exhibited mesenchymal-like traits with FGFR(IIIc) isoforms. By contrast, OSCC cells with low levels of ZEB1/2, and high levels of E-cadherin and ESRP1/2 exhibited epithelial-like traits with FGFR(IIIb) isoforms, similar to the luminal-like subtype of breast cancer[6]. Among the four FGFR paralogs, we only detected expression of FGFR1/2 in OSCC cells. Importantly, mesenchymal-like cancer cells expressed FGFR1(IIIc), whereas epithelial-like cancer cells expressed FGFR2(IIIb) (Fig. 2). During TGF-β–induced EMT in NMuMG cells, TGF-β induces ZEB1/2 expression while repressing ESRP1/2 expression, leading to isoform switching from FGFR2(IIIb) to FGFR1(IIIc), but not to FGFR2(IIIc)[6, 16]. When ESRP1 was ectopically overexpressed during TGF-β–induced EMT, FGFR2(IIIb) changed to FGFR1(IIIb)[6]. Taken together, conversion of FGFR2(IIIb) to FGFR1(IIIc) during EMT requires transcriptional regulation and alternative splicing machinery dependent on ESRP1/2. These observations prompted us to investigate the potential involvement of TGF-β that is secreted autonomously by OSCC cells. When endogenous TGF-β was inhibited by a TGF-β receptor inhibitor, FGFR isoform switching and the regulation of *FGFR1* and *FGFR2* mRNA were not significantly altered in OSCC cells, in agreement with our previous observation in breast cancer cells[6]. Conversely, TGF-β treatment also failed to regulate both isoform switching and transcription of FGFR1/2 (data not shown). Therefore, in addition to isoform switching by ESRP1/2-mediated alternative splicing, signaling pathway(s) apart from the TGF-β pathway may be involved in the expression of FGFR1(IIIc) in mesenchymal-like OSCC.

ZEB1 and ZEB2 are known to be extensively upregulated by TGF-β in both normal epithelial cells and cancer cells[18]. ZEB1/2 suppress the expression of ESRP1 by binding to its promoter region, thereby inducing the expression of FGFR(IIIc) isoforms[6] whereas antagonizing FGFR1(IIIc) downregulates ZEB1 expression (Fig. 3). These findings suggest that, during the early stages of cancer, TGF-β accumulates gradually in cancerous tissues[19, 20] where it subsequently induces EMT by inducing ZEB1/2 expression. Once ZEB1/2 expression has been upregulated, the cells will express FGFR(IIIc) isoforms. Soluble factors, such as FGF2, secreted autonomously by cancer cells undergoing EMT, activate FGFR(IIIc) and ERK pathways to sustain high ZEB1/2 levels. Because FGF2 is itself known to be induced by either TGF-β or FGF2[21, 22], a positive feedback loop of autocrine FGF2 signaling can be initiated by TGF-β and sustained by FGF2, thereby maintaining mesenchymal phenotypes. If this positive feedback loop was generated, the cells maintain EMT phenotypes even in the absence of TGF-β. Following that, abundant distribution of FGF2/4 in the cancer microenvironment further stimulates FGFR(IIIc) isoforms in cancer cells undergoing EMT and sustains EMT phenotypes even in the vascular and lymphatic systems. Thus, the addiction of mesenchymal phenotypes of OSCC cells could be switched from the TGF-β axis to the FGF–FGFR axis. Finally, ZEB1 and ZEB2 promote the recruitment of DNA methyltransferase to the promoter region by interacting with each other, resulting in epigenetic regulation of EMT marker genes such as E-cadherin[7].

FGF2 was previously reported to be produced by the cells of primary prostate carcinomas with metastasis, and that FGF2 causes the switching of FGFR isoforms from IIIb to IIIc[23]. Similar to these observations based on prostate cancer, mesenchymal-like OSCC cells also produced soluble factors that activate FGFR1(IIIc) (Figs. 3B and 3C). However, the pathological significance of isoform switching from FGFR2(IIIb) to FGFR1(IIIc) isoform remains unclear, because both FGFR isoforms have almost the same intracellular structure and tyrosine kinase domains. N-cadherin has been reported to interact selectively with FGFR1(IIIc)[24], suggesting that N-cadherin, expressed in mesenchymal-like OSCC cells, can cooperate to transduce unique signals to sustain high-level expression of ZEB1/2.

An antibody that preferentially recognizes only FGFR1(IIIc) was recently developed[25]. However, antibodies that preferentially recognize specific isoforms of other FGFRs have yet to be generated, probably due to the high degree of structural similarity between their extracellular domains. Therefore, diagnosis and therapy, which specifically target the IIIc isoforms, will require the development of methods that can recognize the IIIc isoform proteins specifically, as well as anti-tumor drugs that can target the IIIc isoform proteins specifically.

## Acknowledgments

We would like to thank Dr. T. Shirakihara and Dr. N. D. Sinh for their helpful advice. This work was supported by THE MITSUBISHI FOUNDATION and JSPS KAKENHI Grant Number 18H02969.

## Conflict of interest

The authors declare that they have no conflicts of interest.

## Abbreviation Lists

EMT: epithelial–mesenchymal transition
TGF-β: transforming growth factor-β
qRT-PCR: quantitative RT-PCR
siRNA: short interfering RNA
OSCC: oral squamous cell carcinoma

